# Preferential interactions of a crowder protein with the specific binding site of a native protein complex

**DOI:** 10.1101/2021.08.06.455410

**Authors:** Xu Dong, Ling-Yun Qin, Zhou Gong, Sanbo Qin, Huan-Xiang Zhou, Chun Tang

**Author notes:** To whom correspondence should be addressed (C.T.) or (H.-X. Z.). X.D. and L.Q. contributed equally. **Data deposition:** The assignment for the backbone amide of HPr in the presence of 30% w/v BSA has been deposited in the BMRB with the accession number of 26310.

## Abstract

The crowded cellular environments provide ample opportunities for proteins to interact with bystander macromolecules, yet direct evidence, let alone residue-specific information, for such nonspecific binding is rare. Here, by combining NMR spectroscopy and atomistic modeling, we investigated how crowders influence the association equilibrium and kinetics of two protein partners, EIN and HPr. Ficoll-70 increases the EIN-HPr binding affinity whereas bovine serum albumin (BSA) decreases the affinity. The opposite effects of the two crowders are quantitatively explained by atomistic modeling, which shows that the stabilizing effect of Ficoll-70 arises from volume exclusion favoring the bound state. In contrast, the destabilizing effect of BSA arises from preferential soft interactions with the free state; notably, BSA has favorable electrostatic interactions with positively charged HPr residues within the EIN-binding site. Some of the residues from this site indeed experience significant chemical shift perturbation when titrated with BSA, while the relaxation rates of HPr backbone amides exhibit overall elevation. Furthermore, relaxation dispersion data indicate that Ficoll-70 and BSA both slow down the EIN-HPr association rate, but change the dissociate rate in opposite directions. The observations on kinetics are accounted for by two effects of the crowders: increasing the solution microviscosity and reshaping the EIN-HPr interaction energy surface. The kind of preferential interactions between BSA and HPr that leads to competition with EIN should be prevalent in cellular environments. Our NMR results and atomistic modeling provide benchmarks, at both qualitative and quantitative levels, for the effects of crowded cellular environments on protein-protein specific interactions.

**Significance Statement:** Although nonspecific binding of crowder macromolecules with functional proteins is likely prevalent *in vivo*, direct evidence is rare. Here we present NMR characterizations showing that bovine serum albumin preferentially interacts with a specific binding site on HPr, leading to competition with the latter’s partner EIN. The preferential interactions result in destabilization of the EIN-HPr native complex and speedup of its dissociation, contrary to expectations from excluded-volume and viscosity effects. Atomistic modeling of macromolecular crowding rationalizes the experimental observations, and provides qualitative and quantitative insight into the influences of the crowded cellular environment on protein-protein specific interactions. Our work also has implications for evolution, regarding how nonspecific binding can be either minimized or exploited for gaining new functions.

---

Contrary to the conditions in most *in vitro* studies using purified components, proteins function in crowded cellular environments. During their functional processes, proteins bind with their specific partners, and at the same time, encounter numerous bystander macromolecules with which nonspecific interactions must occur. All such macromolecular crowders exert two effects: presenting volume exclusion and increasing the microviscosity around functional proteins (1). The excluded-volume effect leads to increases in folding and binding stabilities, as seen in many studies using synthetic polymers such as Ficoll-70 as crowding agents (2). The increase in microviscosity leads to a slowdown of all processes involving the transport of functional proteins, including the diffusion-limited association and dissociation of protein partners.

It is now well recognized that, in addition to volume exclusion, macromolecular crowders also form soft interactions with functional proteins (3, 4). For example, the homodimer of the GB1 A34F mutant is stabilized by the protein crowder bovine serum albumin (BSA) and inside cells, owing to electrostatic repulsion, not volume exclusion, of the GB1 monomers by the crowders (5, 6). Soft interactions can also be preferential, i.e., directed to a specific site, including the site for binding a protein with its specific partner (7-10). Although soft interactions between crowders and functional proteins are intrinsically weak, their distinctions from specific binding between protein partners can be blurred (11). Common properties include site selection (i.e., preferential interactions, as just noted), conformational selection (10), and millimolar affinities (7, 9, 10, 12) that are comparable to those of ultraweak but functional complexes (13-15). On the other hand, seemingly non-binding proteins can gain binding specificity with the introduction of just a few point mutations, as illustrated by the point mutation A34F leading to GB1 dimerization (16).

The effect of crowding on protein-protein interactions has been largely assessed from an energetic perspective in a mostly qualitative fashion. NMR spectroscopy is the method of choice to characterize how the crowded cellular environment modulates protein structure and stability (17). Although NMR can probe dynamic properties, including binding kinetics, of proteins, crowding presents technical challenges. In particular, crowders significantly restrict protein rotational freedom, due to increased microviscosity and nonspecific binding, and thereby deteriorate NMR spectra. In some cases, this apparent drawback, i.e., line broadening or peak disappearance, has been used as an indicator for crowding effects (7, 9, 10). However, kinetic properties are best characterized by the observation, not the absence, of NMR signals. One approach to enhance NMR sensitivity is by introducing an ^19^F probe at a specific residue (6, 18), which unfortunately forgoes residue-specific details of protein-crowder and protein-protein interactions.

A more comprehensive understanding of protein-protein interaction kinetics involves the analysis of both the association and dissociation processes (4, 19). Two protein partners usually form an ensemble of encounter complexes before forming the native, stereospecific complex (20-22). In the example of bacterial phosphotransferases N-terminal domain of enzyme I (EIN) and histidine carrier protein (HPr), it has been shown that the transient encounter complexes between these two enzymes are electrostatically stabilized (20, 21, 23). Mutations to charged residues outside the native interface affect the association equilibrium (24). On the other hand, the dissociation process of protein complexes has been less studied experimentally, let alone in a crowded environment. Indeed, if a crowder preferentially interacts with one or both of the partner proteins either in the free or bound state, we expect distinct effects of crowding on the association and dissociation processes.

An integration of computational and experimental approaches affords deeper insight into how crowding affects protein-protein interactions (10, 25-27). In the present study, we performed solution NMR measurements on the association equilibrium and kinetics of EIN and HPr under both diluted and crowded conditions. With atomistic modeling providing both qualitative explanation and quantitative account, we show that polymer and protein crowders exert opposite effects on the equilibrium constant and dissociation rate constant of EIN and HPr. That is, Ficoll-70 stabilizes the EIN-HPr native complex and slows down its dissociation, whereas BSA destabilizes the native complex and speeds up its dissociation. Most importantly, our computation predicts and our NMR data confirm that BSA preferentially interact with the EIN-binding site on HPr, leading to competition between BSA and EIN for binding with HPr.

## Results

### Polymer and protein crowders change the EIN-HPr binding affinity in opposite directions

We performed NMR titration for ^15^N-labeled HPr with unlabeled EIN protein (**Figure S1**). In both dilute and crowded conditions, the chemical shift perturbations (CSPs) have similar magnitudes and profiles along the HPr sequence (**Figure S2**). The same set of HPr residues, including H15, Q21, V23, G39, S43, L50, T52, and L53, experiences the highest CSPs; these residues are all located within or at the periphery of the interface with EIN in the native complex. Moreover, the perturbed peaks shift progressively along the same directions (**Figure S1**), indicating that the same native, stereospecific complex is formed regardless of the solution conditions.

Fitting the CSPs as a function of the EIN concentration yielded the respective dissociation constants (*K*_D_) under the various solution conditions (**Figure 1** and **Table 1**). The addition of 10% w/v Ficoll-70 increases the binding affinity, as the *K*_D_ value decreases from 41 ± 4 µM in the dilute condition to 31 ± 4 µM in the crowded condition. In comparison, the addition of 10% w/v BSA decreases the EIN-HPr binding affinity, as *K*_D_ increases to 65 ± 5 µM. It is interesting that the polymer and protein crowders change the EIN-HPr binding affinity in opposite directions. We will provide an quantitative explanation for the opposite changes below. Moreover, since *K*_D_ is the ratio *k*_d_/*k*_a_, the change in binding affinity can result from the change in either the association rate constant *k*_a_ or dissociation rate constant *k*_d_, or both. The effects of crowding on the kinetic constants are presented next.

**Table 1.**
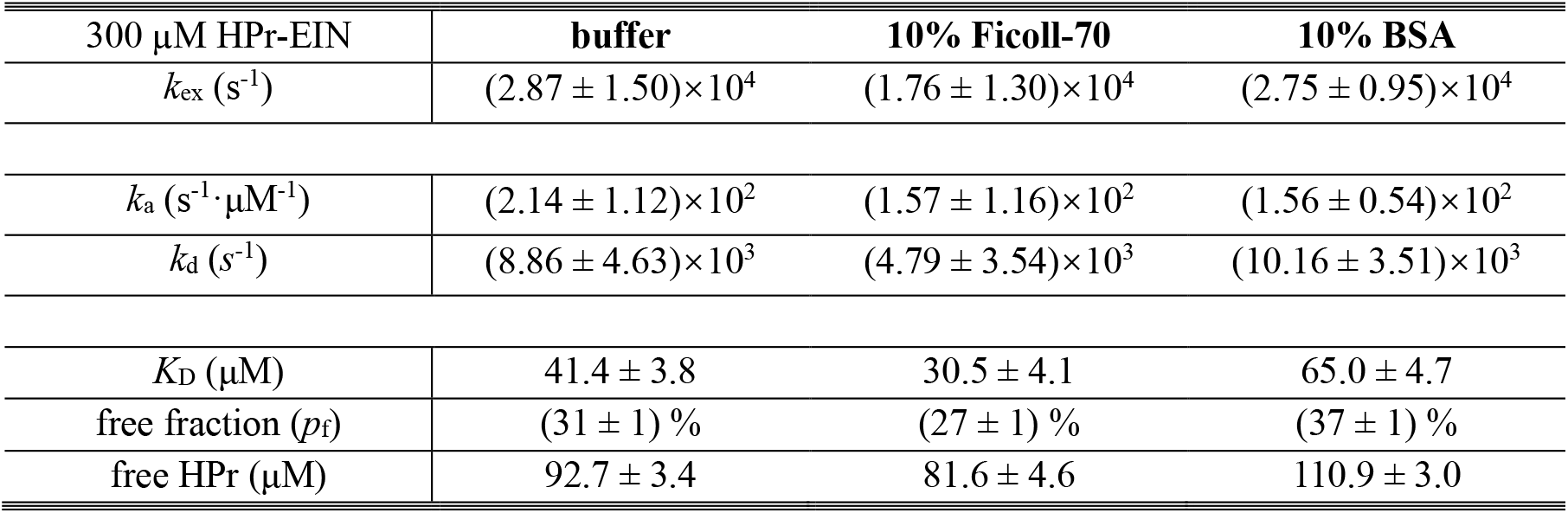
*k*_a_ and *k*_d_ rate constants for EIN-HPr binding from fitting HPr Q51 off-resonance *R*_1ρ_

| 300 μM HPr-EIN | **buffer** | **10% Ficoll-70** | **10% BSA** |
| --- | --- | --- | --- |
| $k_{ex}$ ($s^{-1}$) | $(2.87 \pm 1.50) \times 10^4$ | $(1.76 \pm 1.30) \times 10^4$ | $(2.75 \pm 0.95) \times 10^4$ |
| $k_a$ ($s^{-1} \cdot \mu M^{-1}$) | $(2.14 \pm 1.12) \times 10^2$ | $(1.57 \pm 1.16) \times 10^2$ | $(1.56 \pm 0.54) \times 10^2$ |
| $k_d$ ($s^{-1}$) | $(8.86 \pm 4.63) \times 10^3$ | $(4.79 \pm 3.54) \times 10^3$ | $(10.16 \pm 3.51) \times 10^3$ |
| $K_D$ ($\mu M$) | $41.4 \pm 3.8$ | $30.5 \pm 4.1$ | $65.0 \pm 4.7$ |
| free fraction ($p_f$) | $(31 \pm 1) \%$ | $(27 \pm 1) \%$ | $(37 \pm 1) \%$ |
| free HPr ($\mu M$) | $92.7 \pm 3.4$ | $81.6 \pm 4.6$ | $110.9 \pm 3.0$ |

**Figure 1.**
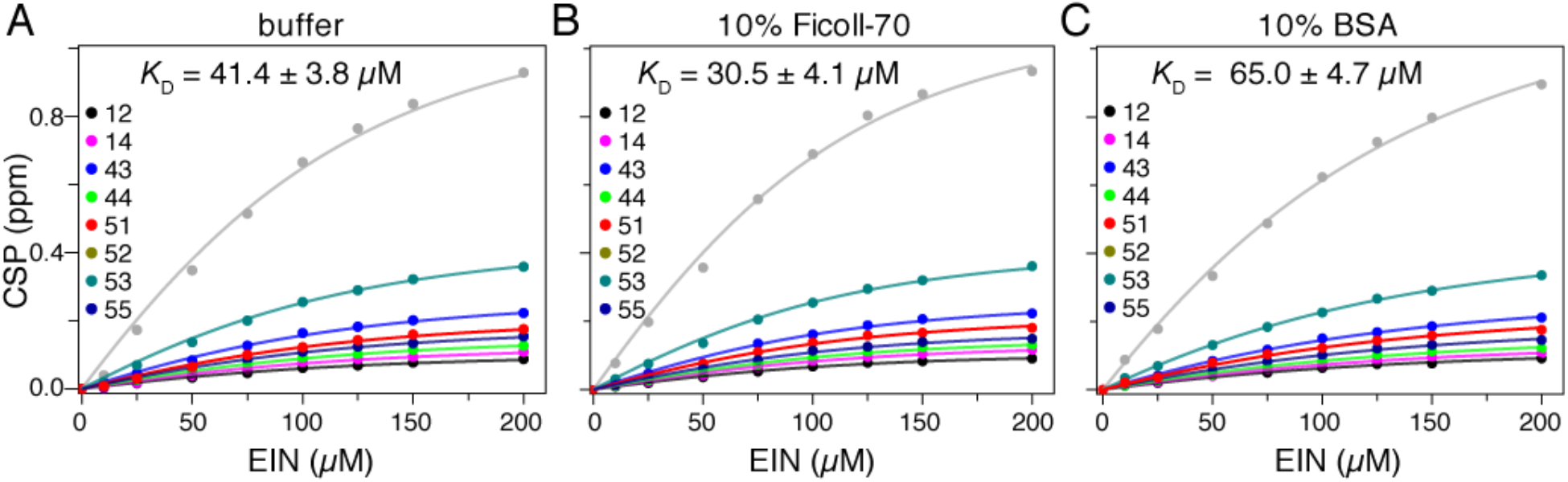
Global fitting of HPr CSPs as a function of EIN concentration. CSPs of 100 µM ^15^N-labled HPr were obtained in (A) the buffer (20 mM Tris•HCl at pH 7.4 with 150 mM NaCl), (B) 10% w/v Ficoll-70, or (C) 10% w/v BSA. The residues included in the global fits and the resulting *K*_D_ values are shown in the legend.

### Crowders modulate kinetic constants but preserve the association and dissociation pathways

To assess how macromolecular crowding impacts the protein-protein association and dissociation kinetics, we performed relaxation dispersion measurements for [^2^H,^15^N]-labeled HPr in the presence of equimolar unlabeled EIN. At an equal concentration of 300 µM, HPr is not fully saturated with EIN and rapidly exchanges between the free and bound states. For diffusion-limited association, *k*_a_ can reach 10^8^ M^-1^•s^-1^ or 100 µM^-1^•s^-1^ (19), and the exchange rate *k*_ex_ = *k*_a_[HPr]_f_ + *k*_d_, where [HPr]_f_ denotes the free HPr concentration, can exceed 10^4^ s^-1^, well beyond the timescale that can be probed by spin-echo relaxation dispersion of ^15^N nuclei (28, 29). Thus, we resorted to the off-resonance *R*_1ρ_ experiment for ^1^H^N^ amide protons. The use of perdeuterated HPr reduces relaxation mechanisms for the protons, while a constant tilt angle during spin-lock minimizes the artifact from the chemical shift differences between the amide protons (30).

We fit the dispersion curves collected from residue Q51 of HPr at two magnetic fields to a two-state fast-exchange model (Eq [2] in Methods) (**Figure 2**). The resulting exchange rates between free and bound HPr are indeed in the order of tens of thousands per second. Data from two other HPr residues, K40 and T59, were noisier but yielded *k*_ex_ values within errors of the values from Q51. Combined with the *K*_D_ values from CSP fitting, we obtained the association and dissociation rate constants (**Table 1**). The association rate constant between EIN and HPr is ∼214 µM^-1^•s^-1^ under the dilute condition, characteristic of a diffusion-limited process accelerated by electrostatic attraction (19, 31). This experimental value is closely matched by a predicted *k*_a_ value of 182 µM^-1^•s^-1^ at the experimental temperature and ionic strength, using our previously developed TransComp webserver (https://pipe.rcc.fsu.edu/transcomp/) (31).

**Figure 2.**
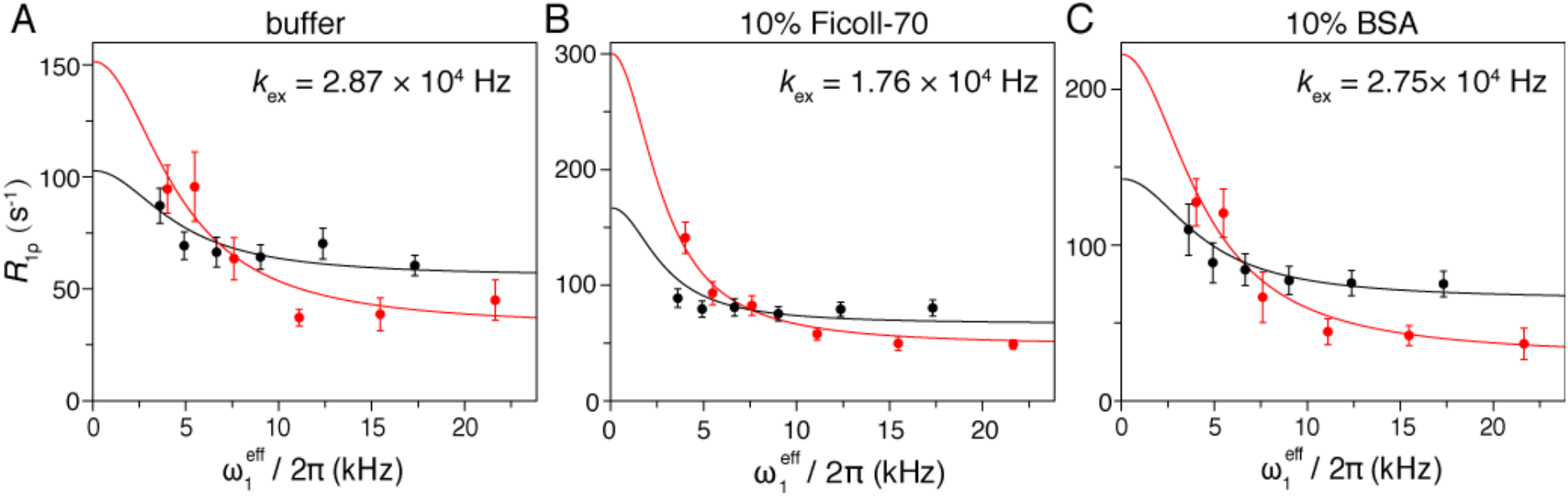
Off-resonance *R*_1ρ_ rates measured for the Q51 amide protons of 300 µM HPr in complex with equimolar EIN at increasing spin-lock field strengths. The experiments were performed in (A) the buffer, (B) 10% w/v Ficoll-70, or (C) 10% w/v BSA, at two magnetic fields (red, 950 MHz; black, 600 MHz). The *k*_ex_ value from a global fit of the two relaxation dispersion curves is shown in the legend. The error bar corresponds to the standard deviation from the *R*_1ρ_ exponential fitting at each spin-lock field.

In the TransComp method, the two protein partners are envisioned to reach a late on-pathway intermediate called the transient complex by translational and rotational diffusion. Moreover, once reaching the transient complex, the two partners rapidly form stereospecific interactions to become the native complex. Hence the entire process is limited by diffusion and the association rate constant is predicted as

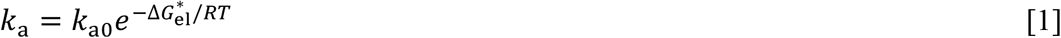

where *k*_a0_ denotes the basal rate constant, i.e., the rate constant by which the two proteins reach the transient complex by free diffusion, and the Boltzmann factor accounts for the acceleration of the diffusion by electrostatic attraction. For EIN and HPr, the basal rate constant (at *T* = 313 K) is calculated to be 1.25 µM^-1^•s^-1^, typical of diffusion-limited association rate constants without electrostatic enhancement (31). The average electrostatic interaction energy, 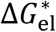, in the transient complex is -3.1 kcal/mol, leading to an enhancement of 146-fold. The strong attraction arises from the significant electrostatic complementarity: the native interface is highly negatively charged on the EIN side but highly positively charged on the HPr side (**Figure 3A**). More specifically, the EIN-binding site on HPr comprises a hydrophobic core (including A20, V23, L47, F48, and L50) and a positively charged rim (including R17, K24, K27, K45, and K49) (**Figure 3B**). Note that the basal rate constant and hence *k*_a_ are proportional to the relative diffusion constant of the two proteins; the latter in turn is inversely proportional to the microviscosity of the solution. A main, but not the only, mechanism by which crowding affects *k*_a_ and *k*_d_ is to increase viscosity.

**Figure 3.**
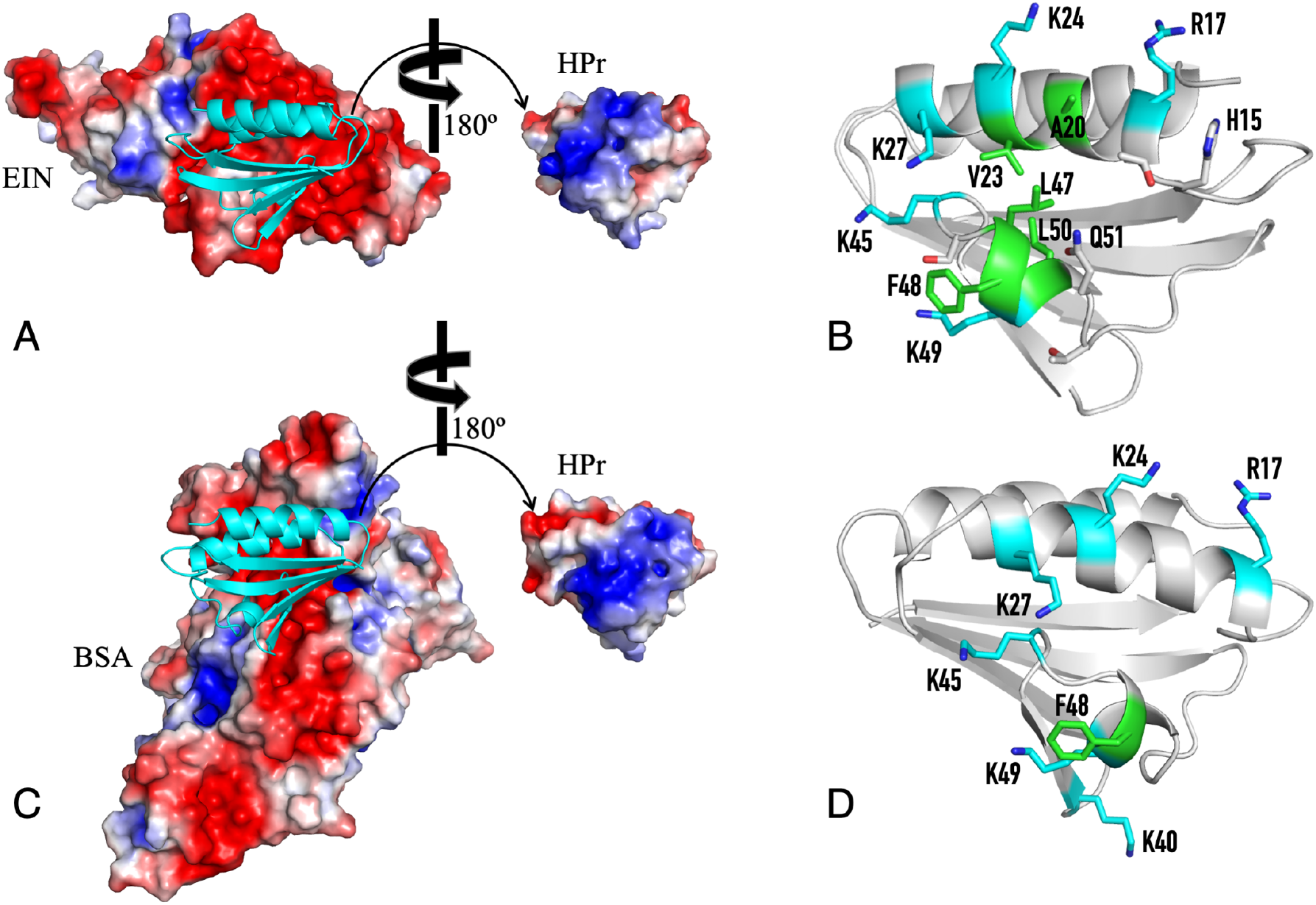
Overlap between the specific binding site of EIN and the preferential interaction site of BSA on HPr. (A) The native complex between EIN and HPr (PDB entry 3EZA). Left: EIN is represented by its molecular surface colored according to electrostatic potential (red: positive; blue: negative), and HPr is shown in cartoon representation in cyan. Right: HPr, rotated 180° to expose the EIN-binding site, is rendered by its electrostatic potential. (B) The EIN-binding site in HPr, with a hydrophobic core (side chains shown as stick with carbon atoms in green) and a positively charged rim (side chains shown as stick with carbon atoms in cyan). (C) A representative complex between BSA and HPr showing significant nonspecific interactions. The HPr molecule has an orientation similar to that in (A), and is presented both as cartoon on the left and electrostatic potential (with 180° rotation) on the right. BSA is presented as electrostatic potential. (D) Residues in free HPr that make the largest contributions to the interaction energy with BSA (c.f. Figure 5B).

Under crowding by both 10% Ficoll-70 and 10% BSA, the association rates deduced from *K*_D_ and *k*ex measurements are slowed by ∼25% (**Table 1**), qualitatively in line with increased viscosity. On the other hand, the dissociation rate decreases by nearly 50% in Ficoll-70 while increases modestly in BSA. Thus, the polymer and protein crowders exert opposite effects on the dissociation process of the EIN-HPr complex, parallel to the corresponding effects on the dissociation constant. A full quantitative account of how crowding affects *k*_a_ and *k*_d_ is given below.

An important question is whether crowding perturbs the association and dissociation pathways. To address this question, we collected intermolecular paramagnetic relaxation enhancement (PRE, denoted as *Γ*_2_) values for the backbone amide protons of EIN. Intermolecular PREs not only report on the native complex, and also probe transient encounters between the two partner proteins (20, 21, 23, 32). As presented above (**Figure S1**), the two crowders do not perturb the native complex of EIN and HPr. With an EDTA-Mn^2+^ probe conjugated at the E5C, E25C, or E66C site of HPr, EIN residues within or near the specific interface, namely residues 52-98 and 103-135, show large PREs under both the dilute and crowded conditions (**Figure 4**). However, for a number of these residues (e.g., residues 78-89 when probed by the HPr E5C mutant and residues 105-132 when probed by the HPr E32C or E66C mutant), the measured PREs are larger than those calculated from the structure of the native complex. The larger-than-calculated PREs are indicative of transient protein-protein encounters (21). Importantly, for all the residues with large PREs, the differences in the PRE values between the crowded and dilute conditions are small (**Figure S3A**), especially when assessed in relative terms (**Figure S3B**). Together, the CSP and PRE data indicate that the macromolecular crowders do not perturb the native or encounter complexes, or the overall pathways for association and dissociation.

**Figure 4.**
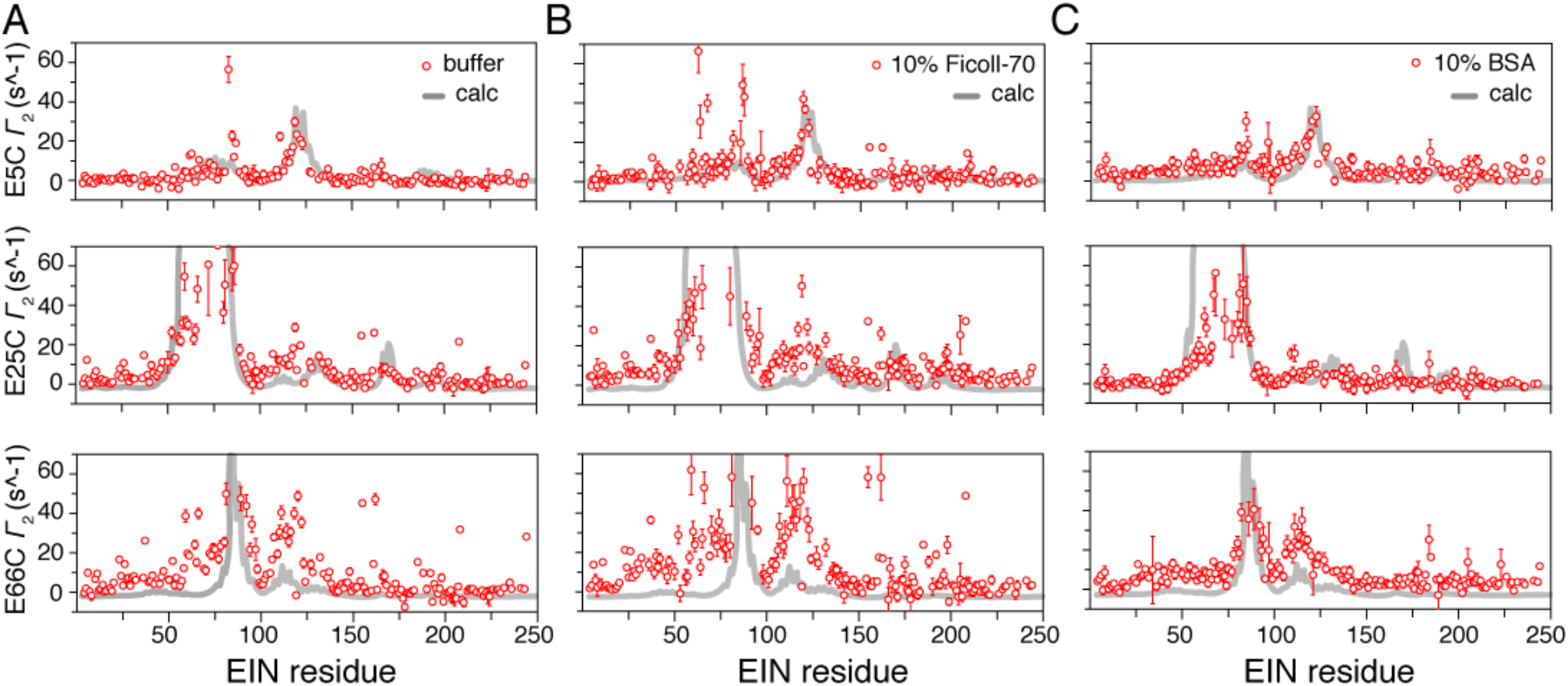
Intermolecular PRE values for backbone ^1^H_N_ protons of 300 µM [^2^H, ^15^N]-labeled EIN, mixed with equimolar unlabeled HPr conjugated with EDTA-Mn^2+^ at E5C (top), E25C (middle), or E66C (bottom). The PRE data were collected in (A) the buffer, or (B) 10% w/v Ficoll-70, or (C) 10% w/v BSA. The error bar of the PRE value corresponds to the standard deviation from the experimental measurement of ^1^H_N_ *Γ*_2_ value using a delay scheme of three time points.

### Volume exclusion dominates Ficoll-70’s interactions with EIN and HPr whereas soft attraction dominates BSA’s interactions with the partner proteins

The increase in EIN-HPr binding affinity by Ficoll-70 is qualitatively consistent with the excluded-volume effect of this crowder (2). We have developed methods to quantitatively account for crowding effects on protein-protein binding affinities (33-35). These methods treat the binding proteins at the atomic level, and compute the change, 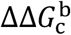, in binding free energy by the crowder (**Figure 5A**). Ficoll-70 is a highly branched, globular polymer that experiences compression as its concentration increases (36). For simplicity, we modeled Ficoll-70 at 10% w/v as hard spheres with a 32-Å radius (estimated by assuming close packing at the solubility limit of 55% w/v). The change in binding free energy by 10% Ficoll-70 is calculated to be - 0.21 kcal/mol (34); the negative value arises because the two proteins present a smaller excluded volume in the bound state than in the free state. This 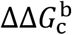 translates to a 1.4-fold increase in EIN-HPr binding affinity, which matches the measured 1.4-fold decrease in *K*_D_, from 41 ± 4 µM in the buffer to 31 ± 4 µM in 10% Ficoll-70, and confirms that the volume exclusion dominates Ficoll-70’s interactions with EIN and HPr.

**Figure 5.**
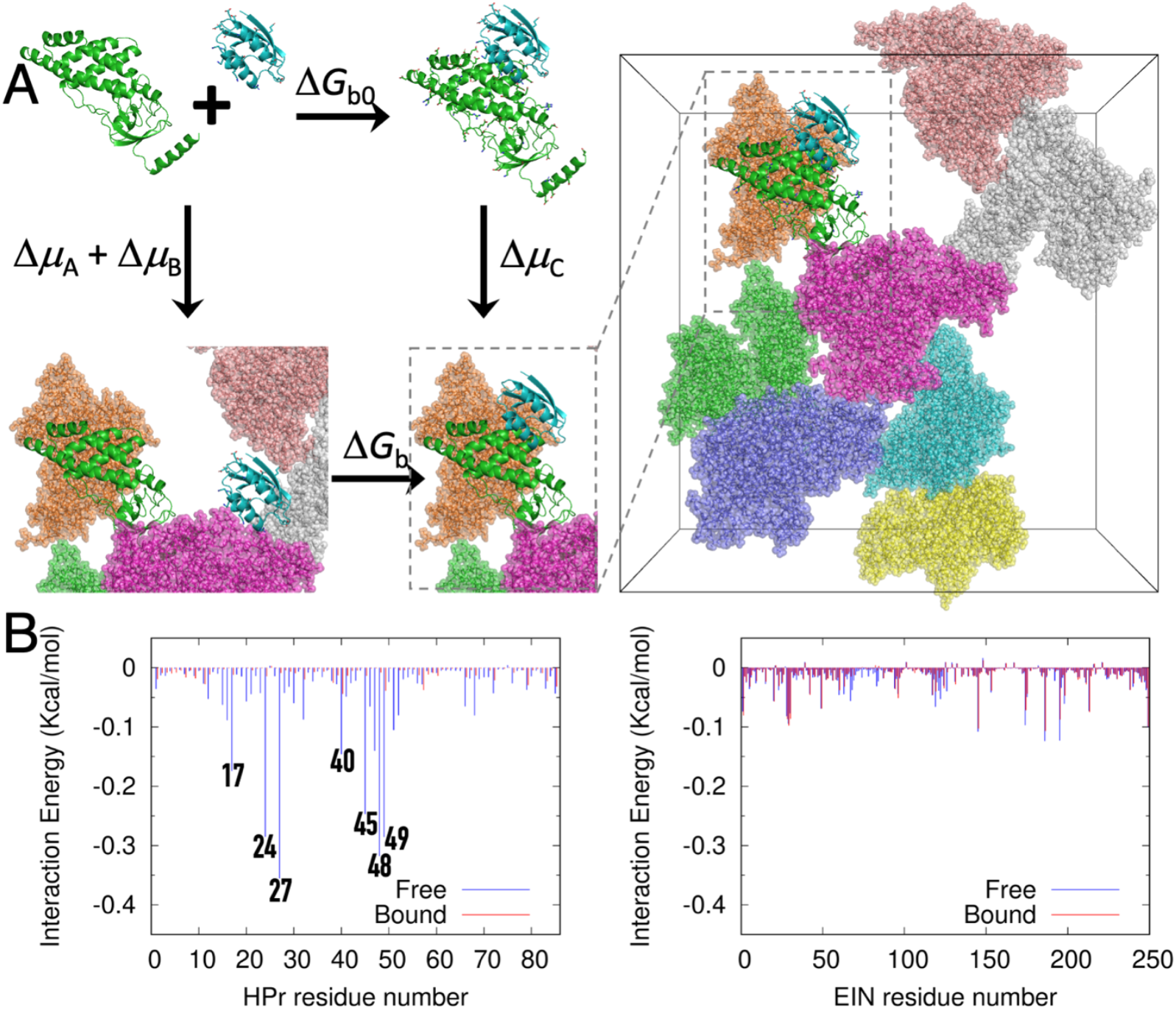
Interactions between the protein crowder BSA and the partner proteins EIN and HPr. (A) Scheme illustrating how the crowding effect of BSA on the binding free energy of EIN and HPr is accounted for. The binding free energies in the dilute and crowded conditions are Δ*G*_b0_ and Δ*G*_b_, respectively. Instead of calculating Δ*G*_b0_ and Δ*G*_b_ (horizontal paths) to find the difference 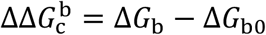, in the FMAP method (35) one calculates the transfer free energies of the two proteins in the free and bound states, denoted as Δ*μ*_A_ + Δ*μ*_B_ and Δ*μ*_C_, respectively, from the dilute to the crowded solution (vertical paths). Because the horizontal and vertical paths close a thermodynamic cycle, one has 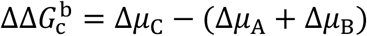. The right panel displays the full view of the EIN:HPr native complex in a box of BSA crowders. (B) Residue-level decomposition of the interaction energies of HPr and EIN with BSA, in either the free or bound state.

For BSA as crowder, we used a more sophisticated of the computational method, FMAP, which treats the crowder also at the atomic level and employs fast Fourier transform (FFT) (35). Now we found 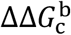 to be +0.33 kcal/mol; the positive value is primarily due to favorable electrostatic interactions between BSA and free HPr (**Figure 3C**). The positively charged surface of HPr complements well the mostly negatively charged surface of BSA. The resulting stabilization of the free state leads to a 1.7-fold decrease in the EIN-HPr binding affinity. This computed crowding effect also agrees well with the measured 1.6-fold increase in *K*_D_, from 41 ± 4 µM in the buffer to 65 ± 5 µM in 10% w/v BSA. If the soft interactions were turned off, the calculated 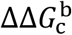 would be -0.26 kcal/mol, similar to that found for Ficoll-70. Therefore, the opposite effects of Ficoll-70 and BSA on the EIN-HPr binding affinity arises because volume exclusion dominates in the interactions between the polymer crowder and the partner proteins whereas soft attraction dominates in the interactions between the protein crowder and the partner proteins.

It is important to recognize that BSA and EIN compete for interacting with the same specific site on HPr, as illustrated in **Figure 3**. Indeed, if BSA were to preferentially interact with HPr sites away from the native interface with EIN, the same preferential interactions would form both in the free and in the bound state, and therefore make no net contribution to 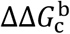. Preferential interactions of BSA can contribute to 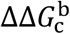 only when they can form in the free but not the bound state, or vice versa.

To confirm that BSA preferentially interacts with the EIN-binding site on HPr, we used a variant of FMAP, FMAPB23 (37), to determine the residue-level decompositions of the BSA-HPr and BSA-EIN interaction energies in the free and bound state. As the results in **Figure 5B** show, the greatest contributions to the interaction energies with BSA come from HPr residues within (or close to) the native interface with EIN, including R17, K24, K27, K40, K45, F48, and K49, in the free state (**Figure 3D**). The interactions of these residues with BSA are prevented in the bound state as they are buried in the interface with EIN. In contrast, individual contributions of EIN residues to the interaction energies with BSA are more modest (**Figure 5B, right panel**). The largest, arising from residues outside the interface with HPr, are nearly the same whether in the free or bound state, and thus have little net effect on 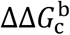. These calculated results will be assessed below against NMR data.

### Crowders affect k_a_ and k_d_ by increasing viscosity and reshaping the interaction energy surface

As illustrated by Eq [1], the association rate constant is determined by the microviscosity of the solution (factored in the basal rate constant) and the long-range interaction energy between the partner proteins (31, 38). Short-range interactions (e.g., van der Waals interactions) help define the location of the transient complex but does not contribute to the Boltzmann factor in Eq [1]. Crowding increases the microviscosity but also reshapes the interaction energy surface (39, 40). If the crowding-induced change, ΔΔ*G*_c_, in interaction energy (which becomes 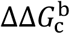 when the partner proteins are in the bound state) is smooth over a ∼10 Å range in the separation between the two partner proteins, then this energetic contribution should be included in the Boltzmann factor (41). Conversely, if ΔΔ*G*_c_ is rugged over this range of protein-protein separation, then this contribution should not enter the Boltzmann factor. For the BSA crowder, the crowding-induced change in EIN-HPr interaction energy is rugged when BSA is treated fully with both volume exclusion and soft interactions (**Figure S4A**). Therefore, the energetic effect of the BSA crowder should have no significant role in changing *k*_a_, leaving the viscosity effect as the major factor. Because the energetic effect applies fully to *K*_D_, which is the ratio *k*_d_/*k*_a_, both the viscosity effect and the full energetic effect of BSA should apply to *k*_d_. When we turned off the soft interactions, the change in EIN-HPr interaction energy due to volume exclusion by BSA becomes smooth (**Figure S4B**), in line with our previous studies (27, 40). As such, for the volume-exclusion crowder Ficoll-70, the viscosity effect and the full energetic effect apply to *k*_a_, but only the viscosity effect applies to *k*_d_.

The microviscosity of Ficoll-70 solutions has been measured by Rashid et al. (42), and at 10% w/v is 2.4-fold higher than the viscosity of water. The viscosity increase by Ficoll-70 thus should reduce both *k*_a_ and *k*_d_ by 2.4-fold. In addition, as presented in the preceding subsection, Ficoll-70 has a 1.4-fold energetic effect favoring association. Therefore, the net effect of 10% w/v Ficoll-70 is predicted to slow down the association rate by 1.7-fold (due to partial cancelation of the viscosity and energetic effects), and slow down the dissociation rate by 2.4-fold (due solely to the viscosity effect). These predictions are in reasonable agreement with measured effects of Ficoll-70 on *k*_a_ and *k*_d_: a 1.4-fold decrease in *k*_a_ and a 1.8-fold decrease in *k*_d_ (**Table 1**).

The bulk viscosity of 10% w/v BSA is 1.6-fold that of water (43), though the microviscosity could differ slightly. As explained above, this viscosity effect is the major factor in the change in *k*_a_ by BSA. This protein crowder also has a 1.7-fold energetic effect favoring dissociation. So the net effects of BSA are predicted to be a 1.6-fold slowdown in association and a 1.1-fold speedup in dissociation. These predictions again are in good agreement with the measured results: a 1.4-fold decrease in *k*_a_ and a 1.1-fold increase in *k*_d_ (**Table 1**).

### Experimental evidence for BSA’s preferential interactions with HPr

One way to probe the interactions between a crowder and a protein is by measuring the relaxation rates of backbone amide nitrogens on the protein (12). In the presence of 10% w/v BSA, the transverse relaxation rate *R*_2_ of free HPr and its product with the longitudinal rate *R*_1_ increase significantly (**Figure 6A**). In contrast, much smaller increases in *R*_1_ × *R*_2_ are observed in the presence of 10% w/v Ficoll-70. Moreover, the addition of either the polymer or the protein crowder causes little increases in *R*_1_ and *R*_2_ for the backbone amide nitrogens of EIN (**Figure S5**). Thus, the protein crowder BSA weakly binds with and thereby slows down the rotational tumbling of HPr.

**Figure 6.**
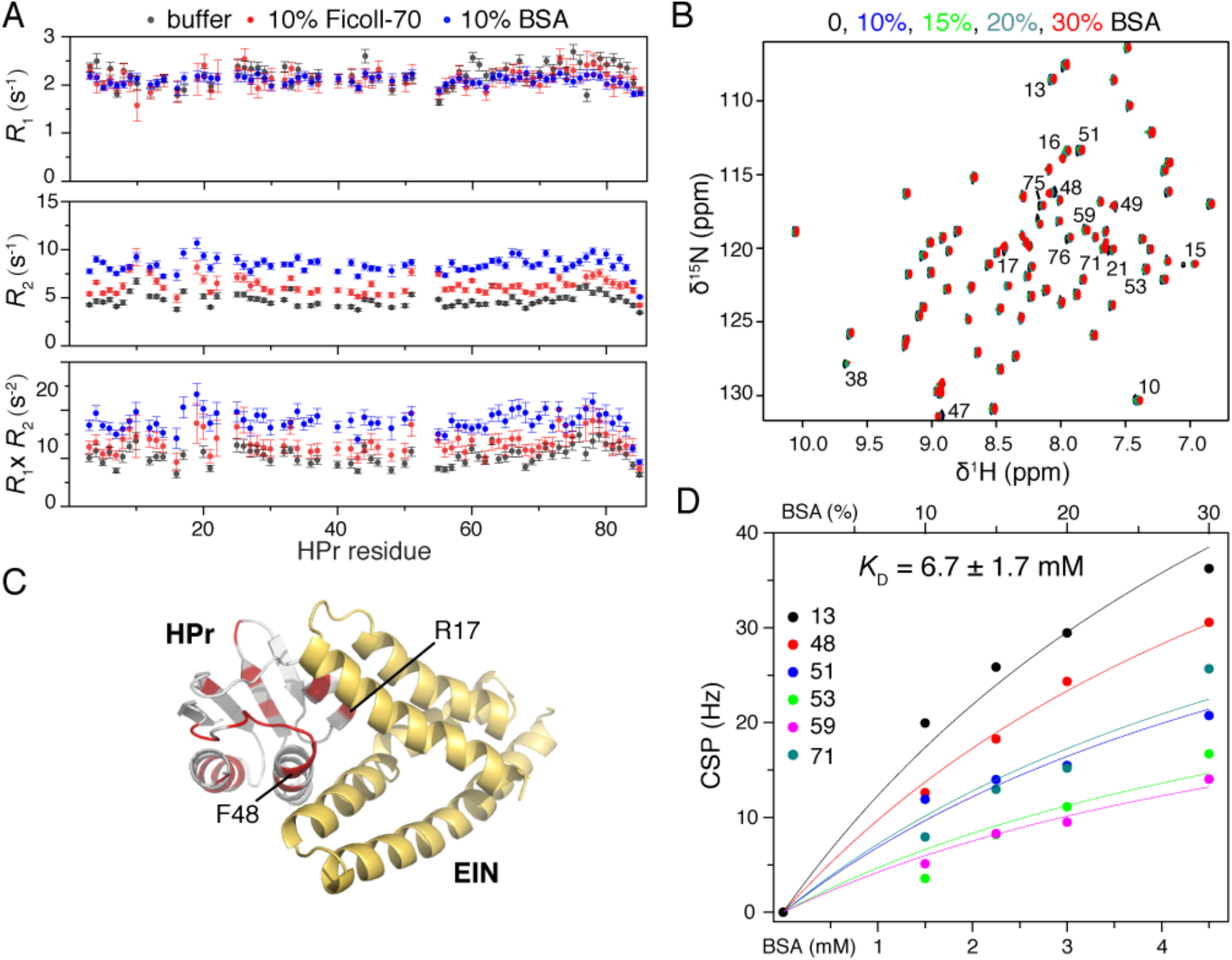
NMR probe of the soft interactions between the protein crowder BSA and free HPr. (A) Longitudinal relaxation rates *R*_1_ (top), transverse relaxation rates *R*_2_ (middle), and the product of *R*_1_ and *R*_2_ rates (bottom) measured for the amide nitrogens of the ^15^N-labeled free HPr, in three solution conditions. The error bar corresponds to the standard devotion from the fitting to the experimental measurement values (top two panels), and the propagated standard deviation (lower panel) (B) Overlay of the NMR HSQC spectra of [^2^H,^15^N]-labeled HPr in the buffer (black), or in the presence of increasing concentrations of BSA. (C) The CSPs obtained at 30% w/v BSA, mapped to the structure of HPr in the native complex with EIN (PDB code 3EZA, colored red), with key interacting residues denoted. (D) The CSPs for six HPr residues are globally fit to a binding isotherm, yielding the BSA-HPr dissociation constant shown in the legend.

We further assessed the changes in *R*_1_ and *R*_2_ for the amide nitrogens of HPr in complex with equimolar EIN. In the presence of 10% BSA, the changes of the relaxation rates are much smaller for EIN-bound HPr than for free HPr (**Figure S6**). These results indicate that BSA-HPr nonspecific interactions are largely abolished upon the formation of the EIN-HPr native complex, which provides additional support for the overlapping interactions sites of EIN and BSA on HPr. This observation also confirms the computational results presented above (see **Figures 3** and **5B**).

To elucidate more structural details about the interactions between BSA and HPr, we carried out NMR titrations for HPr with the protein crowder. With increasing concentrations (from 10% to 30% w/v) of BSA, we observed significant CSPs for a subset of HPr residues (**Figure 6B**). A number of the perturbed HPr residues are located within or near the specific interface with EIN (**Figure 6C**), as predicted by our computation (**Figure 3D** and **Figure 4B, left panel**). Specifically, the perturbed residues including R17 are a part of the positively charged rim of the EIN-binding site on HPr, while the perturbed residues around F48 are a part of the hydrophobic core of the EIN-binding site (**Figure 6C**). As such, BSA and EIN compete for a similar binding site on HPr. Nevertheless, the interaction between BSA and HPr has a millimolar *K*_D_ value, much weaker than that between EIN and HPr (**Figure 6D**). Millimolar crowder-protein interactions have also been reported (7, 9, 10, 12), which are comparable to some specific but ultraweak protein-protein interactions (14, 15, 44).

## Discussion

We have combined NMR spectroscopy and atomistic modeling to study the effects of two crowders, Ficoll-70 and BSA, on the association equilibrium and kinetics of EIN and HPr. The most important observation is that BSA preferentially interacts with the EIN-binding site on HPr, leading to a competition between BSA and EIN for the same binding site. This observation is supported by several lines of evidence, including the increase in transverse relaxation rates of HPr backbone amides in the presence of BSA and abrogation of this effect when EIN is also present, and significant CSPs observed on HPr residues within the EIN-binding site. The preferential interactions between HPr and BSA have profound consequences, including destabilization of the native complex and speedup of its dissociation, contrary to expectations from excluded-volume and viscosity effects of crowders.

The preferential interactions of BSA with HPr are notable because the interaction site on HPr largely overlaps with the specific site for binding the protein partner EIN. Considering the sheer number of macromolecules in cellular environments, preferential interactions are likely prevalent between crowders and functional proteins. One might expect a reduced likelihood that such preferential interactions involve sites of specific binding between protein partners. However, similar to the overlap on HPr between the preferential interaction site of BSA and the specific binding site of EIN, cytoplasmic components preferentially interact with the substrate recognition site on the Pin1 WW domain (8) and with the specific hydrophobic residues on ubiquitin (7). Likewise, *in vitro* studies have shown that both Ficoll-70 and BSA preferentially interact with ligand-binding cleft of the maltose binding protein (9, 10).

Inspection of the BSA surface and the EIN-binding site on HPr reveals only very common features. BSA has an isoelectric point (pI) around 5.0, and thus carries significant negative charges at neutral pH, as illustrated by a mostly negative electrostatic potential surface (**Figure 3C**). The isoelectric points of proteins in all genomes have a bimodal distribution, with peaks around 5.0 and 9.0 (45). Thus most proteins, like BSA, carry net charges at the physiological pH, for solubility, binding, and function (46). The EIN-binding site on HPr is also typical of many other protein binding sites, with a hydrophobic core and a charged rim (46). Therefore, the kind of preferential interactions, mainly electrostatic in nature, that BSA forms with HPr’s EIN-binding site should be common *in vivo*. We further note that it is only because the preferential interaction site on HPr overlaps with the specific binding site that the preferential interactions of BSA favor free HPr over EIN-bound HPr, thereby exerting a net effect on the binding properties of EIN and HPr.

Whereas the effect of crowding on the dissociation constant is solely explained on energetic ground, for diffusion-limited association and dissociation, the increase in microviscosity presents another mechanism for crowders to affect the kinetic rates. For crowders where volume exclusion dominates the energetic effect, previous studies have shown the cancelation of the energetic and viscosity effects on the association rate (27, 40). Our result for the change in *k*_a_ by Ficoll-70 has a similar explanation, even though the cancelation is incomplete so there is still a modest slowdown in association. On the other hand, our observation for a speedup in EIN-HPr dissociation by BSA appears to be unprecedented, and is a clear indication of the viscosity effect being superseded by the energetic effect.

The two crowders studied here, Ficoll-70 and BSA, as far as EIN and HPr are concerned, represent crowders that are dominated by volume exclusion and soft interactions, respectively. Qualitatively, volume-exclusion crowders are expected to stabilize native complexes, whereas crowders exhibiting soft interactions may destabilize native complexes, provided that the soft interactions with the specific binding sites are favorable. In the presence of volume-exclusion crowders, the association of protein partners is pulled in opposite directions by the energetic and viscosity effects, whereas the dissociation process experiences the viscosity effect. In contrast, when crowders exhibit soft interactions, the dissociation is pulled in opposite directions by the energetic and viscosity effects from macromolecular crowding, while it is the association process that mainly experiences the viscosity effect.

The prevalence of preferential interactions by crowder proteins presents the risk that a few mutations could turn them into detrimental, specific interactions (10). The best example is the E-to-V mutation on the β chain of hemoglobin, which leads to polymerization of hemoglobin molecules and the onset of sickle cell anemia. Similarly, a point mutation A34F turned an otherwise monomeric protein GB1 into a homodimer (16). More recently, by introducing three point mutations to a pre-stabilized TEM1 β-lactamase, Cohen-Khait et al. (47) were able to obtain a dimer between mutant and wild-type TEM1 with micromolar affinity. In other studies, promiscuously or weakly interacting complexes of disordered or structured proteins have been linked with gene dosage toxicity (48, 49). Thus, there may be evolutionary pressure to keep preferential interactions in check. On the other hand, the transition from preferential interactions to specific binding may provide an avenue for proteins to gain new functions.

## Methods

### Protein sample preparation

The genes encoding HPr and EIN (the N-terminal domain of enzyme I including residues 1-249) were cloned to the pET11a vector. Cysteine mutations, including E5C, E25C and E66C of HPr, were introduced using the QuikChange approach. BL21* cells were used for expressing the proteins. Uniformly ^15^N-labeled proteins were prepared in M9 minimum medium, with the ^15^N-NH_4_Cl as the sole nitrogen source. Uniformly [^2^H,^15^N]-labeled proteins were expressed in M9 medium with D-glucose-1,2,3,4,5,6,6-d_7_ as the sole carbon source; cell culture was diluted three times before IPTG induction. The proteins were purified using FFQ anion exchange columns and Superdex-75 gel-filtration columns (Cytiva) in tandem. Purified proteins were verified using SDS gel and electrospray mass spectrometry (Bruker Daltonics).

Purified proteins were buffer-exchanged (Amicon Ultra) into 20 mM Tris•HCl buffer at pH 7.4 containing 150 mM NaCl. For macromolecular crowding, the diluted protein solution was carefully mixed with Ficoll-70 (Sigma-Aldrich, Cat# 72146-89-5) or BSA (Sigma-Aldrich, Cat# 9048-46-8) at desired weight/volume (w/v) ratios. The cysteine mutants of HPr were prepared in the same buffer with the addition of 2 mM DTT.

### NMR titrations and relaxation measurements

Unless otherwise indicated, NMR spectra were acquired at 313 K on a Bruker 600 MHz (unless otherwise indicated) spectrometer equipped with a cryogenic probe. The NMR buffer was 20 mM Tris•HCl at pH 7.4 containing 150 mM NaCl, with the addition of 10% D_2_O. A series of ^1^H-^15^N HSQC spectra were recorded to monitor the chemical shift perturbation of 100 µM ^15^N-labeled HPr upon titrating with unlabeled EIN, at concentrations of 25, 75, 125, and 200 μM. The protein samples were prepared either in the buffer, or in the presence of 10% w/v macromolecular crowder. The HSQC spectra were processed using NMRPipe (50), and chemical shift perturbations (CSP, as plotted in **Figures 1 and 6**) were calculated as (0.5×(δH)^2^+0.1×(δN)^2^)^0.5^, where δH and δN were the chemical shift changes in the respective dimensions. For the assessment of soft interactions between BSA and HPr, 200 µM [^2^H,^15^N]-labeled HPr was titrated with 10%, 20%, and 30% w/v BSA.

For measuring the relaxation rates of amide nitrogen atoms, the samples included: (1) 300 µM ^15^N-labeled HPr; (2) 300 µM [^2^H,^15^N]-labeled EIN; and (3) 300 µM ^15^N-labeled HPr mixed with equimolar unlabeled EIN. For crowded conditions, 10% w/v Ficoll-70 or 10% w/v BSA was added. The longitudinal relaxation experiments were performed at recovery delays of 0.01, 0.1, 0.2, 0.35, and 0.5 s. The transverse relaxation rates (*R*_2_) were measured with refocusing delays of 16.97, 50.91, 101.82, 152.73, and 203.64 ms. The *R*_1_ and *R*_2_ relaxation rates were obtained from fitting the relative peak intensities with single-exponential decays.

### Measurement of intermolecular PREs

After removing DTT from the protein buffer by desalting, isotopically unlabeled HPr with a cysteine point mutation (E5C, E25C or E66C) was mixed with maleimide-EDTA (Toronto Research Chemicals, Cat# P996250) pre-chelated with Mn^2+^ for 4 hr. The molar ratios of HPr, maleimide-EDTA, and Mn^2+^ were 1:4:8. The paramagnetically conjugated HPr was further purified on a Source-Q anion exchange column, and successful ligation was confirmed by electrospray mass spectrometry.

A TROSY-based pulse scheme was used to measure ^1^H_N_ transverse relaxation rates of EIN (300 µM, [^2^H,^15^N]-labeled) mixed with equimolar unlabeled wild-type HPr (diamagnetic) or paramagnetically labeled HPr, to obtain the paramagnetic relaxation enhancement (PRE; *Γ*_2_), as previously described (51). The relaxation delay times were set at 8, 20, and 32 ms. The PRE measurements were performed in the buffer or in the presence of the macromolecular crowders. The PREs were calculated based on the structure of the EIN-HPr native complex (PDB entry 3EZA) as described previously (20), with the Mn^2+^ electronic relaxation time at 10 ns and the rotational correlation time of the protein complex at 14 ns.

### Measurement of off-resonance *R*_1ρ_ rates

The sample contained 300 µM [^2^H,^15^N]-labeled HPr and 300 µM unlabeled EIN. The off-resonance *R*_1ρ_ rate constants for ^1^H_N_ were measured using a pulse sequence described previously (30). The spectra were recorded on both 600 MHz and 950 MHz spectrometers. A recycling delay of 2 s was used between scans, and 8 transients were recorded per *t*_1_ increment. The carrier of the off-resonance spin-lock field was adjusted relatively to the proton center (8.2 ppm) according to Δ=±√2ω_1_ (the strength of the rf field), while the strength of the spin-lock field (/2*π*) was varied from 3.3 to 17.7 kHz. The peak intensities of the corresponding spectra were extracted for the subsequent fitting at different spin-lock fields. The *R*_1ρ_ data collected at the two magnetic fields were globally fit to the following equation for a two-state fast-exchange model (52):

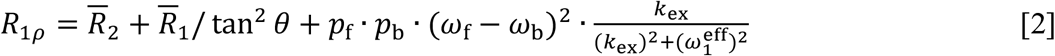

Here 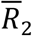 and 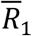 are the relaxation rates in the rotating frame, with the sum, 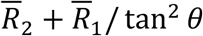, as a fitting parameter; *θ* is the tilt angle of the rotating frame (∼35°); *p*_f_ and *p*_b_ stand for the fractions of free and bound HPr, respectively (calculated from *K*_D_ determined by CSPs as in **Figure 1**); and (ω_f_ – ω_b_) corresponds to the proton chemical shift perturbation upon complex formation (in s^-1^, therefore, the fitted 950 or 600 MHz parameters are related by 2.5-fold). The errors reported in **Table 1** are the standard deviations (and the propagated values) from the fittings.

### Calculation of EIN-HPr association rate constant in dilute condition

The structure of the native complex found in PDB entry 3EZA (53) has the side chain of F48 deeply inserted into crevice on the EIN surface. This F48 conformer led to serious clashes with EIN when we initially applied the TransComp method to determine the transient complex. Because F48 adopts its native conformer only after the formation of the transient complex, we rebuilt the F48 side chain in UCSF Chimera (54), by picking a rotamer (55) that has the side chain retracted from and also avoids clashes with EIN.

The modified PDB file of the complex was uploaded to the pdb portal of the TRANSCOMP web server (https://pipe.rcc.fsu.edu/transcomp/frompdb.html) (31). The server converted the pdb file to pqr format using pdb2pqr (56), with atomic partial charges assigned for pH = 7.4 and atoms assigned the Bondi radii. The server then sent the data to the computer clusters at Florida State University’s Research Computing Center to do the calculations. The calculations involved three steps: determination of the transient complex by mapping the interaction energy surface via sampling the 6-dimensional space of relative translation and relative rotation; finding the basal rate constant by Brownian dynamics simulations; and computing the electrostatic interaction energy in the transient complex by the APBS method (57). Sample results at ionic an strength of 0.17 M and a temperature of 298 K are available at https://pipe.rcc.fsu.edu/transcomp/out/32a396b47d2832507b57017643025bd2/4689/. The final results were corrected to the experimental temperature of 313 K.

### Calculation of the crowder-induced change in EIN-HPr interaction energy

Our FMAP method (<u>F</u>FT-based <u>M</u>odeling of <u>A</u>tomistic <u>P</u>rotein-crowder interactions) was developed to calculate the transfer free energy of a protein from a dilute solution to a crowded solution (35). In this method, protein-crowder interactions (including steric, nonpolar, and electrostatic terms) are expressed as correlation functions, which is a direct product of two functions in Fourier space. The strength factor for the nonpolar term was 0.16; the strength factor for the electrostatic term was 1.6; the cutoff distance was 12 Å. The ion strength was 0.17 M, and the temperature was set to 313 K.

To prepare the crowded solution, BSA was built by homology modeling using a human serum albumin structure (PDB entry 1AO6) and eight copies were placed in a cubic box with a side length of 199.423 Å (concentration at 111 g/L) and simulated in explicit solvent for 70 ns (35). The eight copies of BSA in the final snapshot was saved as representing the crowded solution. For each test protein (e.g., HPr), the Boltzmann factor of the protein-crowder interaction energy was averaged over grid points (334 along each dimension) and over 4392 orientations of the protein, to yield the transfer free energy, Δ*μ*, from a dilute solution to the crowded solution. For each EIN-HPr complex, this calculation was done three times, once for the complex (“C”) and once for each of the two partner proteins (“A” and “B”). The difference, Δ*μ*_C_ − (Δ*μ*_A_ + Δ*μ*_B_), is finally the crowder-induced change in EIN-HPr interaction energy.

This calculation was done for the native complex to yield 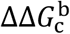, the crowding effect of BSA on the binding free energy, and also when the two partner proteins were separated to a given distance, to yield ΔΔ*G*_c_ as a function of the inter-protein separation. In the latter case, the two partner proteins were separated along the normal direction of the least-squares plane of the interfacial atoms (31). If the resulting configuration had clashes between the partner proteins (which could happen when the separation was < ∼4 Å), we replaced it with the closest one sampled by TransComp. The selection for the closest configuration was based on root-mean-square-deviation.

### Residue-level decomposition of BSA-EIN and BSA-HPr interaction energies

Energy decomposition was carried out by considering the interaction of a test protein, e.g., HPr, with a single BSA molecule (taken as copy #1 in the crowder box), using our FMAPB23 method (37). We selected the 1000 HPr-BSA pair configurations with the lowest interaction energies according to FMAPB23, and then recalculated their interaction energies by the atom-based method (which finds the interaction energy by enumerating all pairs of atoms between the protein-crowder pair). For each HPr-BSA pair configuration, the interactions between all the atom pairs connecting one residue of the protein with one residue of the crowder were added up, and this partial sum was divided by two, with one half assigned to the protein residue and one half assigned to the crowder atom. The total contribution of each protein residue was accumulated over all the crowder residues as interaction partners, and a final average over the 1000 protein-crowder configurations was taken. This calculation was done both on the EIN-HPr native complex, yielding the residue-level decomposition in the bound state, and on the two partner proteins, yielding the decomposition in the free state.

## Acknowledgement

We thank the Beijing NMR Center and the NMR facility of the National Center for Protein Sciences at Peking University for providing technical support and assistance in data collection and analysis. The work has been supported by the National Key R&D Program of China (2018YFA0507700 to C.T. and 2017YFA0505400 to X.D.), by the National Natural Science Foundation of China (21921004 to C.T.), and by the National Institutes of Health (Grant GM118091 to H.-X.Z.).

## Notes

### Competing Interest Statement

The authors have declared no competing interest.

