## Supplementary Figures for "Preferential interactions of a crowder protein with the specific binding site of a native protein complex"

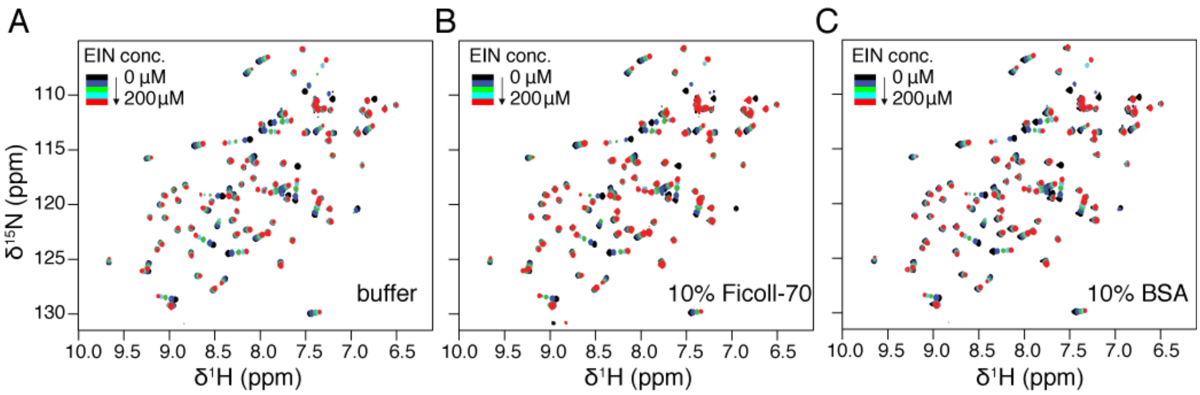


**Figure S1.** Overlay of the HSQC spectra of 100 μM $^{15}$N-labeled HPr at increasing concentrations of unlabeled EIN. The HSQC spectra were collected in (A) the buffer (20 mM Tris•HCl at pH 7.4 with 150 mM NaCl), (B) 10% w/v Ficoll-70, or (C) 10% w/v BSA. The titration of EIN into HPr caused CSPs in the same set of residues along the same directions, regardless of the solution conditions.

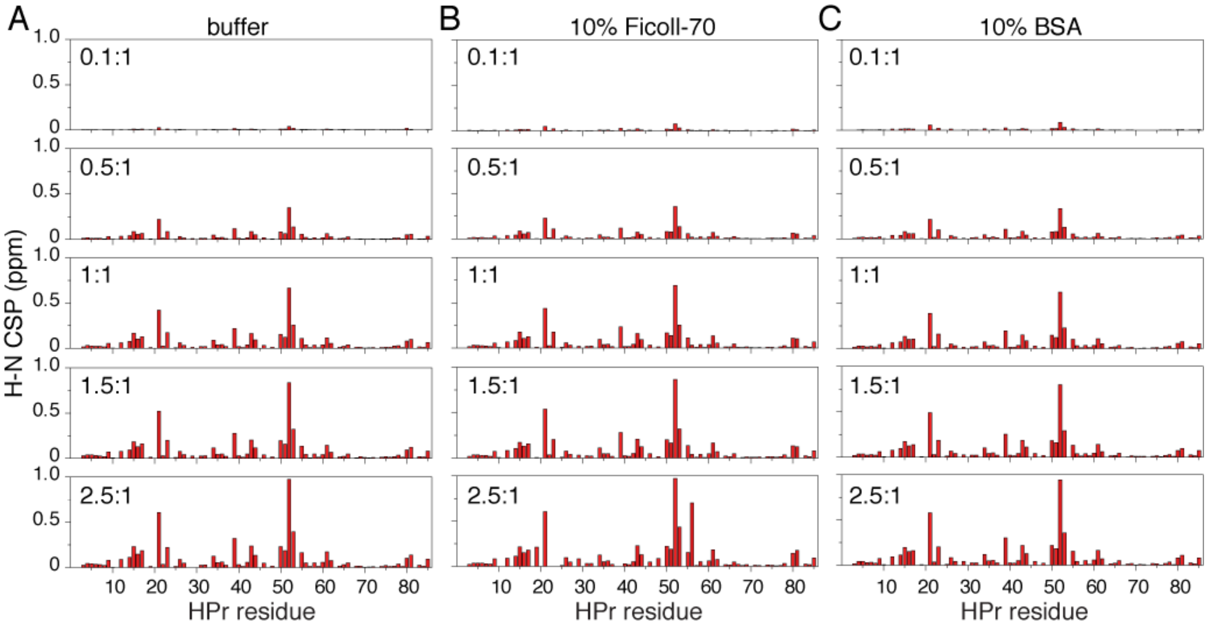


**Figure S2.** CSP profiles for 100 μM $^{15}$N-labeled HPr upon the addition of unlabeled EIN. The CSPs were measured in (A) the buffer, (B) 10% w/v Ficoll-70, or (C) 10% w/v BSA.

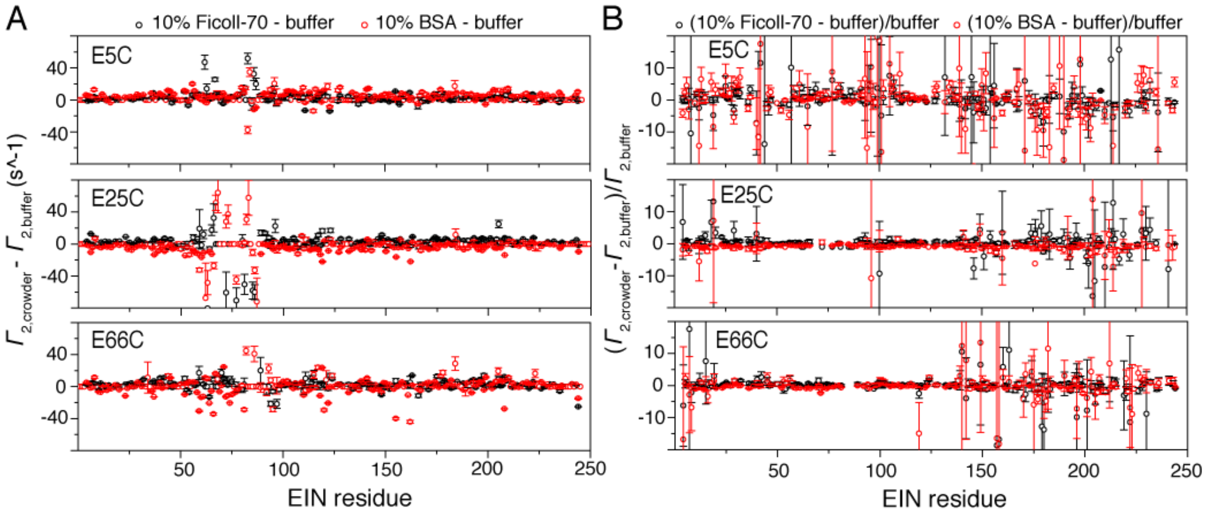


**Figure S3.** The difference in the intermolecular PRE $\Gamma_2$ values for backbone $^1H_N$ collected in the diluted buffer or in the presence of macromolecular crowders. (A) The absolute differences in the intermolecular PRE $\Gamma_2$ values. The PRE values in the crowded conditions are multiplied by a factor (0.9 for Ficoll-70 and 1.14 for BSA) before subtraction, to account for the difference in $K_D$ and bound fraction. (B) The relative PRE differences. For each residue, the absolute PRE difference in (A) is further divided by the PRE value observed in the diluted condition.

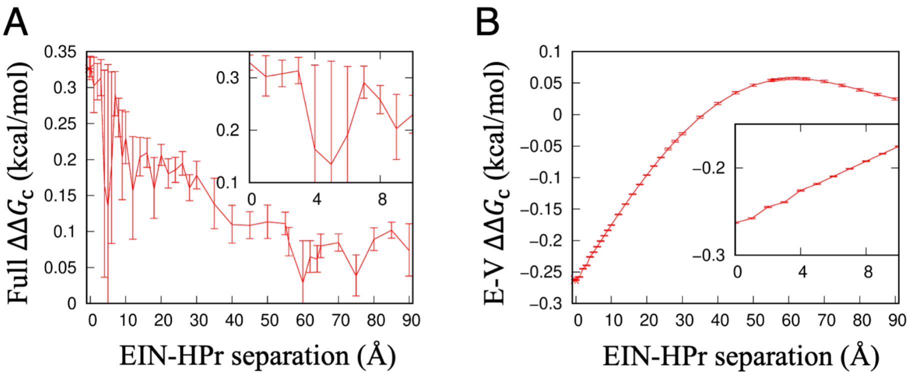


**Figure S4.** The change, $\Delta\Delta G_{\mathrm{c}}$, in the interaction energy between EIN and HPr by the presence of 10% w/v BSA. $\Delta\Delta G_{\mathrm{c}}$ is calculated in the same way as $\Delta\Delta G_{\mathrm{c}}^{\mathrm{b}}$ illustrated in Figure 5A, except that EIN and HPr are placed at various separations. (A) $\Delta\Delta G_{\mathrm{c}}$ results when both volume exclusion by and soft interactions with BSA are fully accounted for. (B) $\Delta\Delta G_{\mathrm{c}}$ when only the excluded-volume ("E-V") effect of BSA is considered. Error bars represent variations among FMAP results obtained from 4392 orientations of the protein complex.

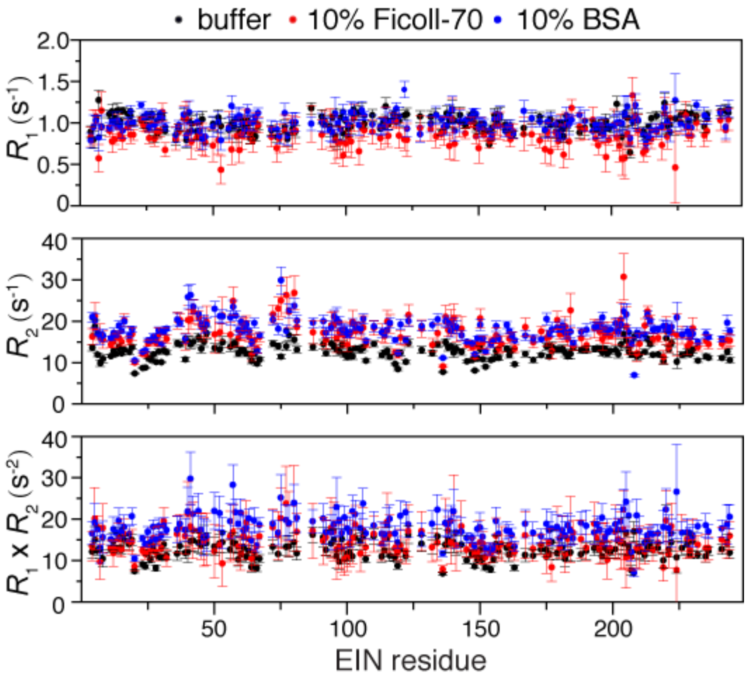


**Figure S5.** NMR probe of the soft interactions between the protein crowder BSA and free EIN. Longitudinal relaxation rates $R_1$ (top), transverse relaxation rates $R_2$ (middle), and the product of $R_1$ and $R_2$ rates (bottom) are measured for the amide nitrogen atoms of [$^2$H,$^{15}$N]-labeled free EIN, in three solutions conditions.

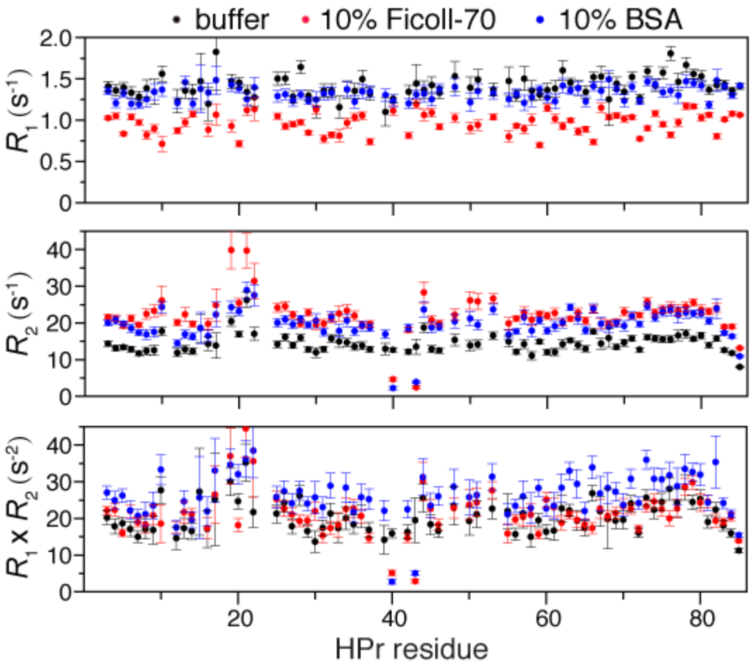


**Figure S6.** NMR probe of the soft interactions between BSA and HPr in complex with EIN. Longitudinal relaxation rates $R_1$ (top), transverse relaxation rates $R_2$ (middle), and the product of $R_1$ and $R_2$ rates (bottom) are measured for the amide nitrogen atoms of [$^2$H,$^{15}$N]-labeled HPr in complex with equimolar unlabeled EIN at 300 μM.
